# in vivo RNA structural probing of guanine and uracil nucleotides in yeast

**DOI:** 10.1101/2022.09.29.510215

**Authors:** Kevin Xiao, Homa Ghalei, Sohail Khoshnevis

**Affiliations:** Department of Chemistry, Emory University; Department of Biochemistry, Emory University School of Medicine

## Abstract

RNAs have critical catalytic or regulatory functions in the cell and play significant roles in many steps of gene expression regulation. RNA structure can be essential for its cellular function. Therefore, methods to investigate the structure of RNA in vivo are of great importance for understanding the role of cellular RNAs. RNA structural probing is an indirect method to probe the three-dimensional structure of RNA by analyzing the reactivity of different nucleotides to chemical modifications. The chemical modifications can target either the RNA backbone or the Watson-Crick face of nucleotides. The selective 2’-hydroxyl acylation analyzed by primer extension (SHAPE) can probe the ribose sugar in all unpaired RNA nucleotides. In contrast, Dimethyl sulfate (DMS) alkylates adenine and cytosine and reports on base pairing context but is not reactive to guanine (G) or uracil (U). Recently, new compounds were used to modify Gs and Us in the plant model system *Oryza sativa* and in the prokaryotic organisms *Bacillus subtilis* and *Echerichia coli*, as well as human cells. To complement the scope of RNA structural probing by chemical modifications in the model organism yeast, we analyzed the effectiveness of guanine modification by a family of aldehyde derivatives, the glyoxal family, in *Saccharomyces cerevisiae* and *Candida albicans*. We also explored the effectiveness of uracil modification by carbodiimide N-cyclohexyl-N-(2-morpholinoethyl) carbodiimide metho-p-toluenesulfonate (CMCT) in vivo. We show that among the glyoxal family, phenylglyoxal (PGO) is the best guanine probe for structural probing in *S. cerevisiae* and *C. albicans*. We also demonstrate uracil modification by CMCT in *S. cerevisiae* in vivo. Further, we show that PGO treatment does not affect the processing of different RNA species in the cell and is not toxic for the cells under the conditions we have established for RNA structural probing. Our results provide the conditions for in vivo probing the reactivity of guanine and uracil in RNA structures in yeast and offer a valuable tool for studying RNA structure and function in two widely used yeast model systems.

## Introduction

RNAs constitute a significant class of nucleic acids that serve essential cellular functions. In eukaryotes, protein-coding messenger RNAs (mRNAs) comprise only ~4% of the total cellular RNAs (1). In contrast, most transcribed eukaryotic RNAs are non-coding and can play critical catalytic or regulatory roles in gene expression. These non-coding RNAs comprise the majority of the total cellular RNAs. As RNAs can be both single- and double-stranded and are highly flexible, they can adopt diverse secondary and tertiary structures in physiological conditions. An example of simple RNA three-dimensional structures includes base-paired doubled-stranded areas such as hairpin stems. Complex RNA three-dimensional structures comprise tertiary and quaternary structures such as ribose zippers, kink turns, and pseudoknots (2). The versatility of RNAs in forming simple and intricate three-dimensional (3D) structures allows RNAs to perform critical cellular functions, including catalysis, and promote RNA-protein interactions.

Understanding the 3D structures of RNAs is essential for revealing RNA functions in gene expression. However, the diverse range of RNA functions happens in the active forms of RNA in vivo. Researchers, therefore, face the challenge of probing a vast array of RNA structures in vivo to elucidate the functional roles of RNAs in cellular processes (6). RNA structure-function relationships inside the cell are affected by the transcription rate, the local solution environment, and the presence of small molecules or RNA-protein interactions. The physical state of RNA in vivo can provide insight into the function of the RNA (7). Structural biology approaches, including X-ray crystallography, NMR spectroscopy, and single-particle cryo-electron microscopy, have shaped much of our understanding of the 3D structures of many RNAs. While providing us with the atomic resolution structure of the RNA molecules, these techniques have limitations, including the need for highly pure samples and the time it takes to solve the structures. Furthermore, these techniques to study the RNA molecules in vitro may not provide a comprehensive view of the conformations that the RNA molecules adopt in their native environment (8). Hence, indirect techniques have been developed as an effective alternative to structural approaches to study the RNA structure in vivo and in vitro.

RNA structure probing is the fastest way to indirectly investigate RNA structures by using chemical reagents to modify nucleotides at specific positions and analyze the modification efficiency of each site. RNA structure probing techniques can target ribose sugars or nitrogen bases. <u>S</u>elective 2’-<u>h</u>ydroxyl <u>a</u>cylation analyzed by <u>p</u>rimer <u>e</u>xtension (SHAPE) uses an electrophilic carbonyl derivative that can be attacked by the strong nucleophile 2’-hydroxyl group on the ribose sugar (9). Modifications in the RNA backbone result in stops during the reverse transcription (RT). Thus, RT stop sites reveal the modification adduct locations. Quantifying these stops by denaturing UREA polyacrylamide gel electrophoresis (UREA-PAGE) or deep-sequencing provides a measure to assess the accessibility and/or conformation of each nucleotide (10). Base-specific modification is an alternative method for indirect RNA structural probing. Dimethyl sulfate (DMS) is used as a base-specific probing reagent against the N1 of adenines and N3 of cytosines that are not involved in a base pairing or hydrogen bonding (11). DMS is cell-permeable because of its small size. Therefore, DMS modification reactions readily occur in nearly all in vivo conditions without additional permeabilization (11). However, DMS is unable to probe uracils or guanines. The lack of reliable chemical probes for guanines and uracils limits the potential of RNA structure probing approaches to assess RNA conformations. Thus, new families of chemicals have been explored to probe guanines and uracils in vivo.

In the glyoxal family, glyoxal (GO), methylglyoxal (MGO), and phenylglyoxal (PGO) are potential candidates for probing guanines. These chemical probes are carbonyl derivates that are electrophilic towards the nucleophilic amidine group in adenine, cytosine, and guanine. As uracil lacks this functional group, it does not react with glyoxal family compounds. However, because guanines have the amidine group farthest away from the ribose sugar backbone, there is less steric hindrance than in adenines and cytosines, so guanine is most reactive with glyoxal family compounds (12). Glyoxal family compounds were shown to be effective guanine probing agents in the eukaryotic model *Oryza sativa* and gram-negative bacteria *Bacillus subtilis* and *Echerichia coli* in vivo (13). The use of glyoxal and its derivatives has not been established for in vivo RNA structure probing in other model systems.

Uracils are probed by carbodiimides family reagents, including 1-ethyl-3-(3-dimethyl aminopropyl) carbodiimide (EDC) and carbodiimide N-cyclohexyl-N-(2-morpholinoethyl) carbodiimide metho-p-toluenesulfonate (CMCT). EDC reaction in vitro is slower than CMCT due to hyperconjugation, weakening the electrophilicity of the compound. Therefore, CMCT is supposed to be a much faster and thus better reagent than EDC for probing uracils due to the strengthening of electrophilicity from the inductive effect of the toluenesulfonate functional group. However, due to the lack of cell permeability of CMCT, EDC was preferred for its rapid cell wall penetration ability and validated for uracil probing in vivo (14,15).

*Saccharomyces cerevisiae* and *Candida albicans* are two well-studied fungi that also serve as model organisms for studies of conserved biological processes. *S. cerevisiae* is a simple eukaryotic organism whose RNA biology shares many features with higher eukaryotes, making it a suitable model organism to study different aspects of gene expression (16). *C. albicans* is an opportunistic fungal pathogen and the most prevalent cause of fungal infections (17). While there are established procedures for in vivo probing of the adenine and cytosine positions within yeast RNAs using DMS (11,18,19), guanines and uracils have so far escaped in vivo probing in yeast cells. In this work, we test the application of glyoxal and carbodiimide derivatives for RNA structure probing in the widely used budding yeast model system *Saccharomyces cerevisiae* and in the human fungal pathogen *Candida albicans*. We compare three different glyoxal derivatives and show that PGO yields the highest modification rate of guanines in yeast without affecting cell viability or RNA processing. We also present the conditions for in vivo modification of uracils in yeast using CMCT.

## Materials and Methods

### Yeast Cell Culture

BY4741 strain *Saccharomyces cerevisiae* and BWP17 strain of *Candida albicans* were grown in YPD media at 30°C to an optical density of 0.5-0.6 before applying the desired chemical probes.

### Chemical Treatment

*S. cerevisiae* and *C. albicans* were incubated with glyoxal family compounds (GO, MGO, PGO) or CMCT. For each compound, three different concentrations were tested. A no-compound solvent-treated sample was used as the negative control. Two incubation times (5 and 15 min) were tested for all conditions, and two biological replicates were analyzed. GO concentrations were 30 mM, 60 mM, and 120 mM. MGO and PGO concentrations were 5 mM, 10 mM, and 20 mM. CMCT concentrations were 25 mM, 50 mM, and 100 mM. All incubations were performed at 30°C. After incubation, samples were cooled down immediately on ice, and cells were harvested by centrifugation. Cells were washed three times with ice-cold water before RNA extraction.

### Total RNA Extraction and Purification

Total RNA was extracted from cell pellets harvested from 10 mL of culture grown to mid-log phase using hot acid phenol. The extracted RNA was treated with DNase I for 15 minutes at 37°C and further purified using a Quick-RNA miniprep kit (Zymo Research) according to the manufacturer’s protocol.

### Reverse Transcription (RT)

Approximately 1 μg of purified RNA in 10 μL was mixed with 2 μL of 0.6 μM ^32^P-labeled primers. Annealing was performed by incubation at 65°C for 5 minutes followed by a gradual cool down at room temperature over 10 minutes and final incubation on ice. The reverse transcription was performed using SuperScript III (ThermoFisher) at 50°C for 5 minutes per the manufacturer’s manual. 1 μL of 4M NaOH was added before heating the RNA at 95°C for 5 minutes to remove the RNA templates. The cDNA products were mixed with the formamide loading dye, heated at 95°C for 5 minutes, and separated on a prewarmed 6% urea/acrylamide sequencing gel in 0.5X TBE buffer. The gel was dried and exposed to a phosphoscreen.

### Northern Blot for snoRNA and tRNA

Total RNA from two biological replicates treated with PGO or CMCT was isolated using the hot phenol method. snoRNAs were separated on 8% acrylamide/urea gels, transferred to Hybond nylon membrane (GE Healthcare), and probed using ^32^P-labeled DNA oligos against U3-3’ (AAAGTGGTTAACTTGTCAG), tRNA^Leu^ (GCATCTTACGATACCTG) and scR1 (ATCCCGGCCGCCTCCATCAC).

### Serial Dilution Spot Test for Chemical Toxicity Assessment

*S. cerevisiae* was incubated with PGO or CMCT as described above. The density of the culture was adjusted to a final concentration of 10^7^ cells/mL, followed by four successive cascade dilutions in a 1:10 ratio. Dilutions were spotted onto YPD plates and grown at 25°C, 30°C, and 37°C for 48 hours.

## Results and Discussion

### PGO is highly effective for probing guanine nucleotides in S. cerevisiae

To define the best glyoxal derivative for probing guanines in yeast, we tested different concentrations of glyoxal (GO), methylglyoxal (MGO), and phenylglyoxal (PGO) at two incubation times. The tested concentrations of GO and MGO were chosen based on their effect on yeast cell growth, as a proxy for their cell penetration (20). Based on this analysis, 60 mM GO and 10 mM MGO resulted in ~ 50% growth rate reduction in *S. cerevisiae* cells. Therefore, we tested these concentrations as well as 0.5X and 2X of each (30, 60, and 120 mM GO, and 5, 10, and 20 mM MGO). PGO concentrations were chosen based on the effect on yeast mitochondrial ATP synthase (21,22), where 10 mM PGO greatly destabilized the F1-ATPase in *S. cerevisiae*. We, therefore, tested 5, 10 and 20 mM concentrations of PGO.

The 5.8S rRNA is a part of the large ribosomal subunit. Several positions on 5.8S are subject to glyoxal modification in rice 5.8S rRNA, including G82, G89 and G99 (equivalents of G78, G85 and G95 in *S. cerevisiae)* (12). To probe the effectiveness of GO, MGO and PGO, we therefore studied the modification of the 5.8 rRNA of *S. cerevisiae* by these compounds (**Figure 1**). GO weakly modifies guanines in the 5.8S rRNA of *S. cerevisiae* in vivo, as evident from the weak modification of nucleotides at positions G78, G85 and G95 (**Figure 1A**). At the 5 min time, GO modifications do not show a noticeable concentration-dependent relationship, as band intensity across all GO concentrations appears similar (**Figure 1A**). At the 15 min time, the band intensity for G78 and G85 increases as the GO concentration increases (**Figure 1A**). The reactivity of MGO towards guanines in the 5.8S rRNA of *S. cerevisiae* in vivo is weaker than that of GO (**Figure 1B**). G78, G85, and G95 modifications are comparable across all MGO concentrations, as band intensity remains the same at each time point and at each MGO concentration (**Figure 1B**). In contrast, PGO demonstrates the high reactivity towards guanine nucleotides in the 5.8S rRNA of *S. cerevisiae* in vivo (**Figure 1C**). Further, at G78 and G85, modifications by PGO appear to be concentration-dependent, as band intensity at both positions increases as the PGO concentration rises (**Figure 1C**).

**FIGURE 1.**
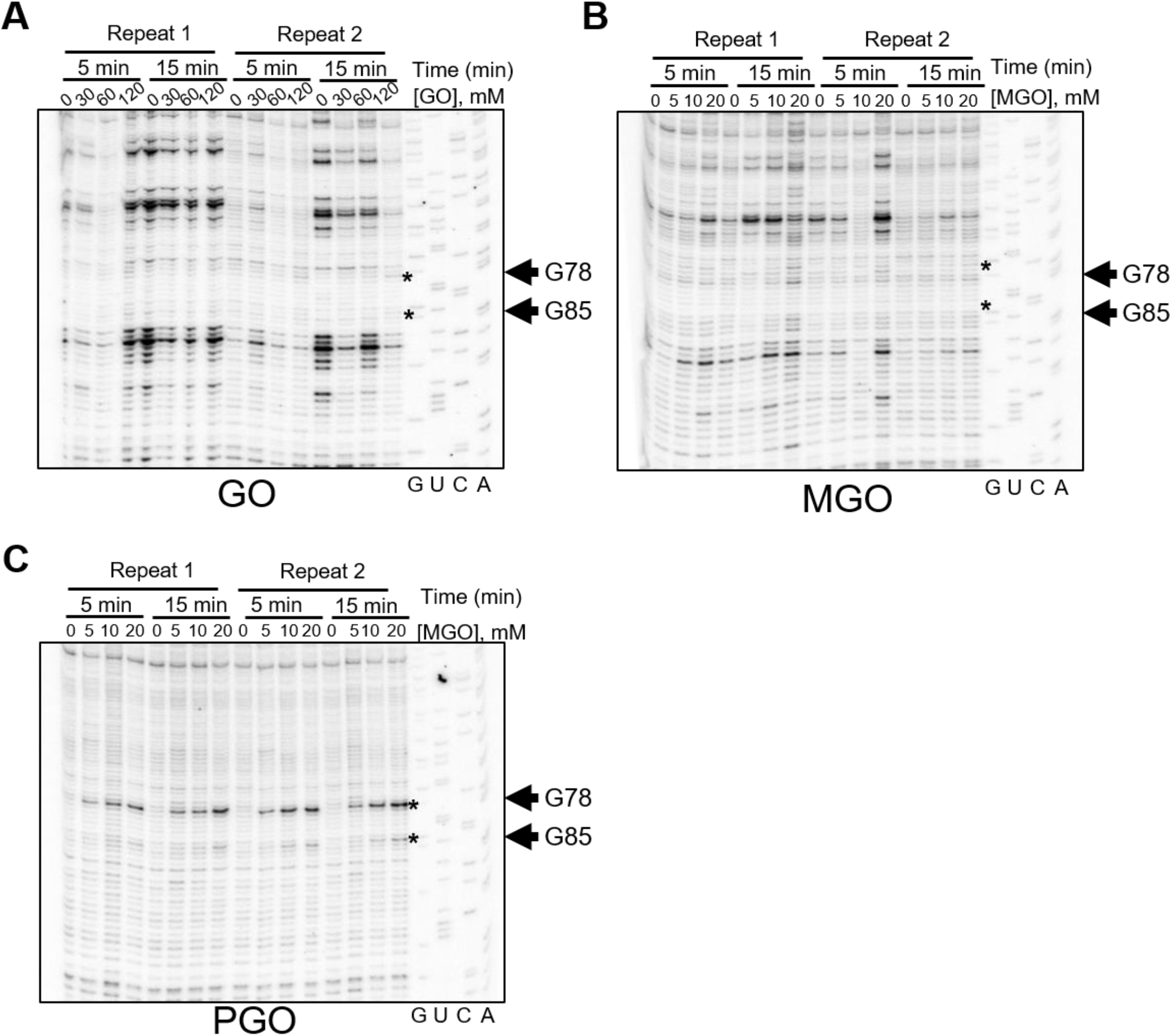
In vivo modification of yeast *S. cerevisiae* 5.8S rRNA by glyoxal and derivatives characterized by denaturing UREA-PAGE. RNA structure probing with **(A)** glyoxal **(B)** methylglyoxal, and **(C)** phenylglyoxal. Reaction conditions are specified at the top of each panel. Two biological replicates were tested in each experiment. Arrows and the asterisks indicate RNA nucleotide positions modified by glyoxal derivatives.

Unlike in rice, PGO does not modify the G95 in *S. cerevisiae* 5.8S rRNA. To understand the reason for this, we analyzed the position of 5.8S G95 in the structure of yeast ribosomes (23). In this structure, G95 is engulfed by the C-terminal tail of the ribosomal protein RPL37. Particularly, residues Gln79 and Ser82 of RPL37 sandwich the base of G95 and come in hydrogen-bonding distance with the N1 of G95, thereby stabilizing its protonated state. Thus, G95 in *S. cerevisiae* 5.8S is sequestered by the C-terminal tail of RPL37 providing an explanation for the lack of reactivity of this nucleotide with PGO (**Figure S1**). The high reactivity of PGO comes from its increased hydrophobicity conferred by the phenyl function group in the molecule, which allows it to penetrate through the phospholipid membrane bilayer (12). The hydrophobic phenyl group on PGO can strengthen interactions between PGO and hydrophobic protein residues, orienting the electrophilic PGO carbonyl group in place for nucleophilic attack by the amidine of guanine (12).

### PGO treatment does not cause RNA processing defects

A critical concern in RNA structural probing is whether the RNA metabolic pathways in the cell changed upon treatment with the chemical probing reagents. We therefore assessed how different RNA processing pathways are affected by PGO treatment by looking at the processing of small nucleolar RNA U3 (U3 snoRNA) and a transfer RNA (leucine tRNA) using Northern blot analysis (**Figure 2A**). Total RNA extracted from yeast cells treated with various concentrations of PGO at two time points was used for the Northern blot analysis. We first analyzed the processing of U3 snoRNA, required for proper ribosome biogenesis. U3 is transcribed as a precursor with 5’- and 3’-extensions which need to be removed in order to form the mature U3 snoRNA (24). We used probes against sequences within the 3’ region of the U3 precursor or within the scR1 RNA as the loading control. The band intensities for U3-3’ species remain the same relative to the loading control (scR1 RNA) even after the concentration of PGO increases, indicating that the treatment of yeast cells with PGO does not impact the processing pathway of U3 snoRNA (**Figure 2B**). As another control, we analyzed the processing pathway of the leucine tRNA (tRNA^Leu^) which is transcribed as a precursor and undergoes processing before entering the translation pool (25). We probed the mature and precursor tRNA using an oligo complementary to the mature part of the tRNA. Our analysis indicates that the levels of mature and precursor of tRNA^Leu^ do not change in the presence of PGO relative to the loading control (scR1) (**Figure 2B**). These data indicate that the tRNA^Leu^ processing pathway remains unchanged in the presence of PGO. Together, our data indicate that treatment of yeast cells with PGO under our established conditions for RNA chemical probing are unlikely to affect the processing of major RNA species in the cell.

**FIGURE 2.**
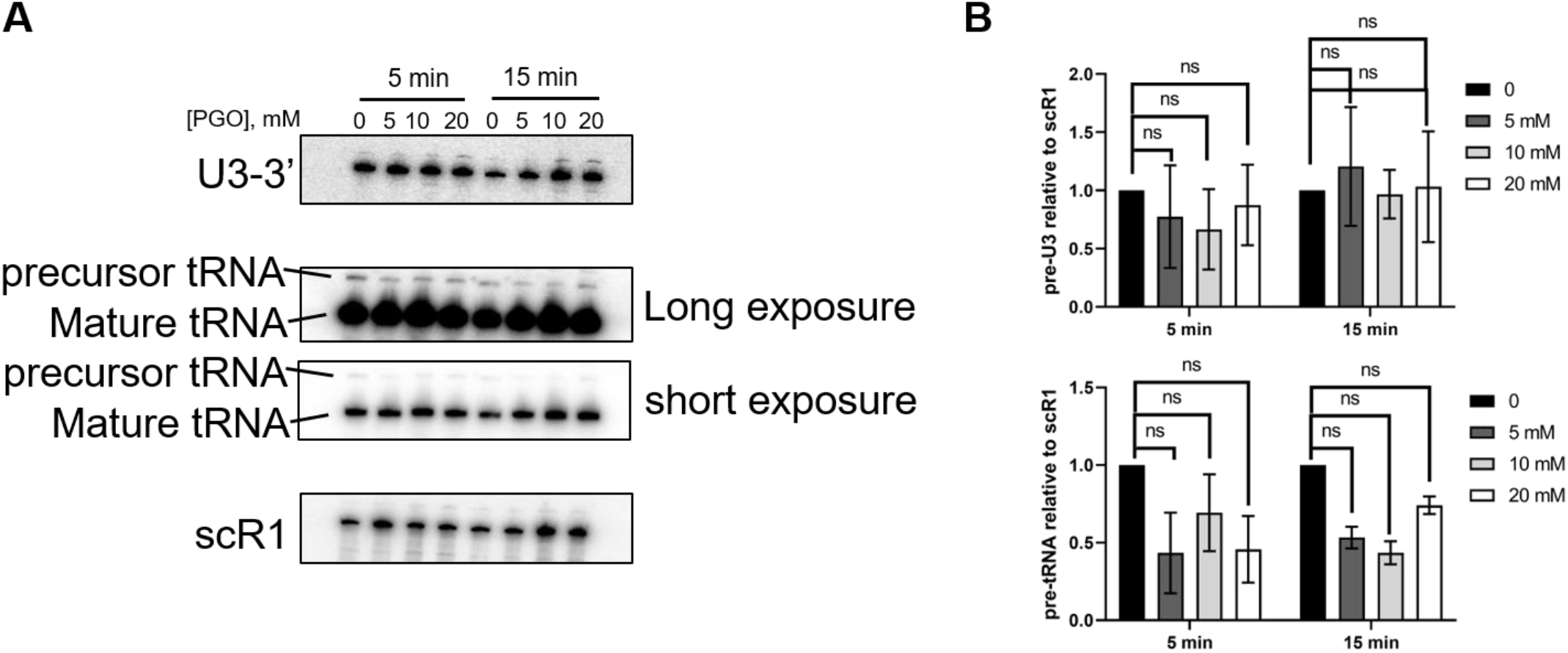
PGO treatment does not affect the processing of U3 and tRNA^Leu^ RNAs in *S. cerevisiae*. **(A)** Northern blot analysis of the U3 and tRNA^Leu^ processing. Reactions were performed at two specified timepoints and three concentrations of PGO (5, 10, 20 mM). scR1 RNA served as the loading control. **(B)** quantification of the data in panel A. Graphs represent two independent experiments. Significance was determined using a t test; n.s., nonsignificant.

### Long treatment with high PGO concentration affects cell growth in S. cerevisiae

As there are other pathways which can affect different RNA molecules in the cell and are not studied here, we analyzed the overall fitness of yeast cells upon PGO treatment. A serial dilution spot test was conducted on PGO-treated *S. cerevisiae* (**Figure 3**). At concentrations below 20 mM, PGO does not cause toxicity for *S. cerevisiae* as judged by the similar size of individual colonies of treated cells compared to no PGO-treated cells (**Figure 3**). However, at the concentration of 20 mM and 15-minute incubation period, *S. cerevisiae* cells no longer withstand the toxicity of PGO, resulting in smaller colonies and slow growth (**Figure 3**). Although this can be due to the toxic effect of PGO on the mitochondrial ATP synthase (21,22), we cannot rule out the possible effects of PGO on other RNA processing pathways in yeast. Based on these data, 5-minute incubation of *S. cerevisiae* cells with 20 mM PGO appears to be the optimal condition to achieve efficient nucleotide modification without affecting the cell viability.

**FIGURE 3.**
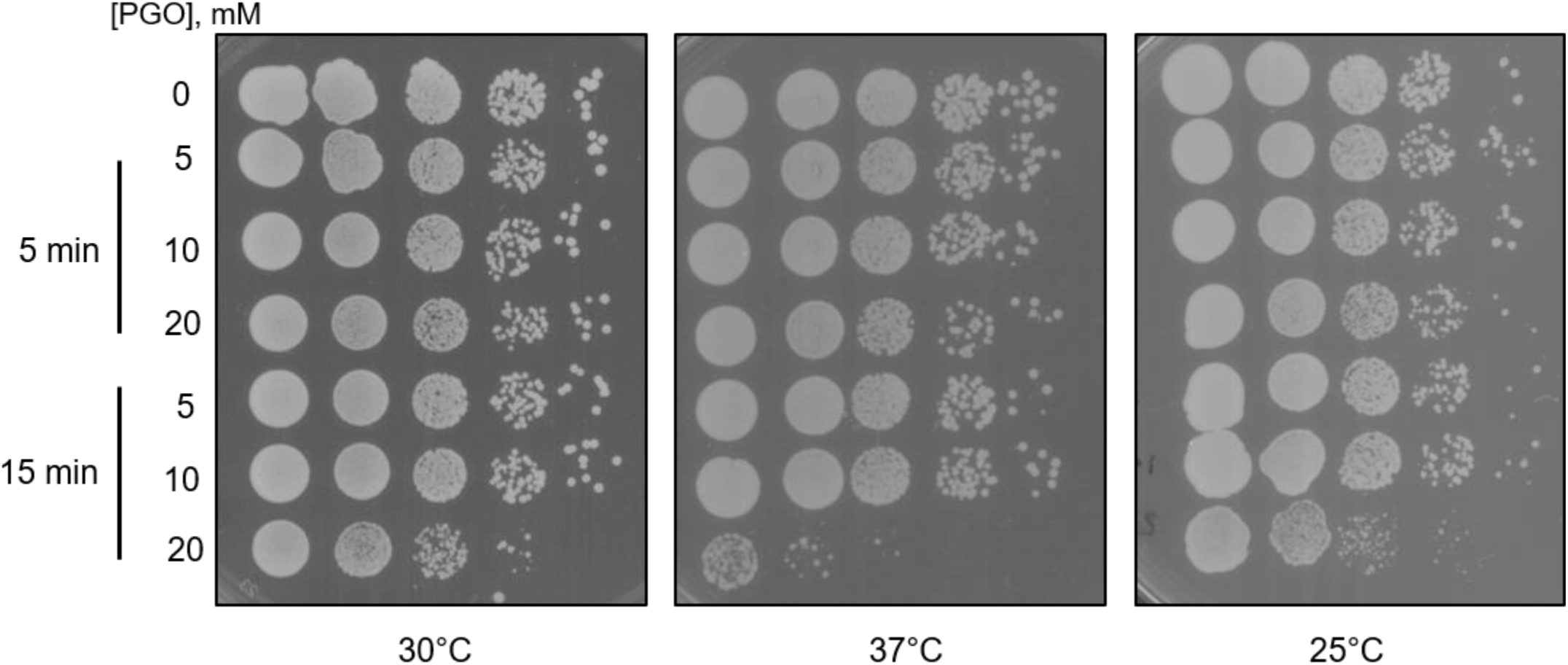
Serial dilution test for toxicity analysis in PGO-treated *S. cerevisiae*. Reaction conditions at two timepoints and concentrations of PGO are specified at the left side of the figure. Plates were incubated at 30°C, 37°C, and 25°C as indicated.

### PGO can be used to modify Gs in other yeasts

Next, we sought to determine whether the conditions for modifying guanines in *S. cerevisiae* are applicable to other fungi. To this end, we tested the modification of 5.8S rRNA of *Candida albicans*, a human fungal pathogen. Post-transcriptional regulation of gene expression is important for the pathogenicity of *C. albicans* (26). Given the emergence of the human fungal pathogen *C. albicans* as a public health threat, it is important to develop tools to study gene expression in this organism. Therefore, establishing the effective condition for guanine probing in *C. albicans* is of great importance for future pharmaceutical and biochemical applications. Having already established that PGO is the best guanine probe in *S. cerevisiae*, we used PGO to probe the 5.8S rRNA in *C. albicans*. PGO shows effective modification reactivity towards G77 and G84 in *C. albicans* (equivalent to G78 and G85 in *S. cerevisiae)* (**Figure 4**). PGO modification of guanine demonstrates weak band intensities overall, compared to that in *S. cerevisiae*, representing lower frequency of guanine modifications in *C. albicans* than that in *S. cerevisiae*. To understand the basis for this difference, we used the published structures of the 80S ribosome from *S. cerevisiae* (23) and *C. albicans* (27) to compare the 5.8 rRNA in these organisms. In both organisms, G78 (G77 in *C. albicans)* is flanked between A77 and A79. However, A79 is flipped out in *C. albicans* 5.8S (**Figure S2**). As a result, while the N1 and N2 of G78 in *S. cerevisiae* face the N1 and C2 of A79, in *C. albicans*, the N1 and N2 of G77 face the N6 and N7 of A78, respectively. We speculate that the different nucleotide configuration in *C. albicans* results in the reduced activity of the N1 of G77, causing the lower reactivity of this nucleotide against PGO in *C. albicans* compared to *S. cerevisiae*. An alternative explanation could be that PGO entry to *C. albicans* cells is less efficeint compared to *S. cerevisiae*. Together, our data suggest that PGO can be used as a suitable chemical reagent for probing guanines in *C. albicans*.

**FIGURE 4.**
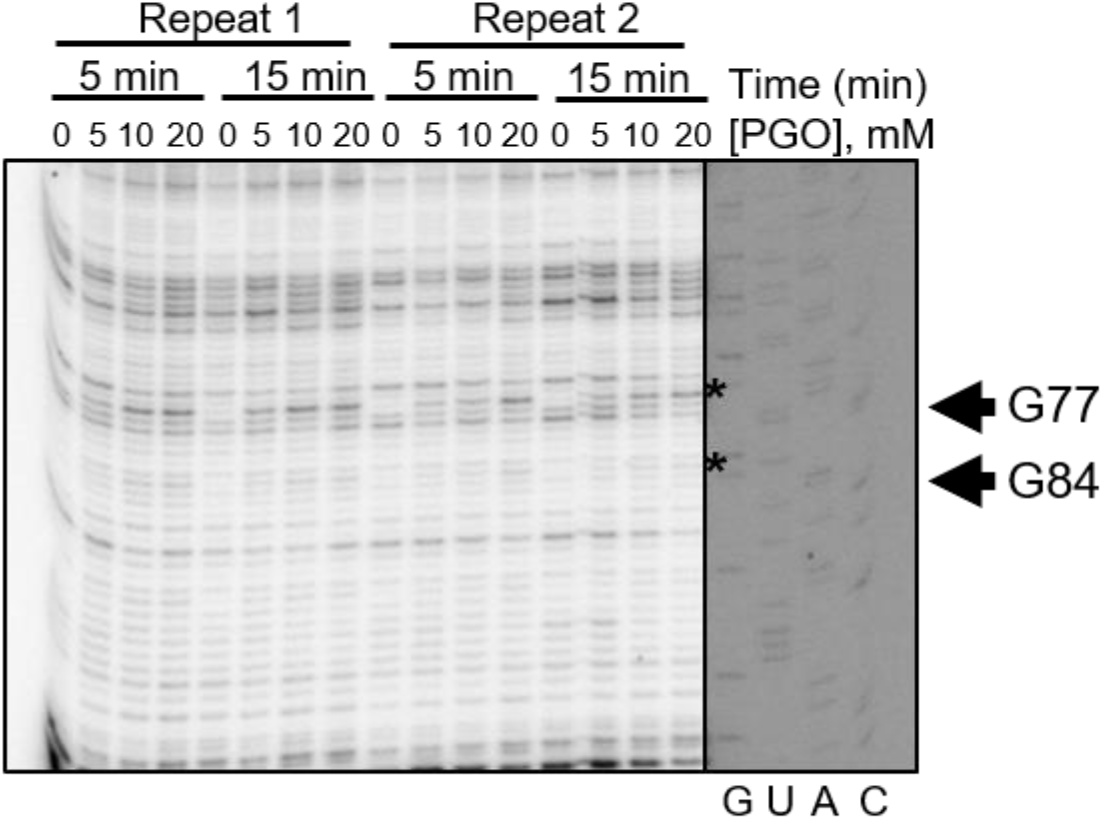
In vivo modification of *C. albicans* 5.8S rRNA by phenylglyoxal analyzed by denaturing UREA-PAGE. Reaction conditions at two timepoints and three PGO concentrations are specified above the gel. Arrows represent modified RNA nucleotide positions by PGO.

### CMCT can be used to modify uridines in S. cerevisiae

Uracil probing chemical reagents and their reaction conditions have yet to be established in yeast in vivo. Carbodiimide N-cyclohexyl-N-(2-morpholinoethyl) carbodiimide metho-p-toluenesulfonate (CMCT), a carbodiimide derivative, has been used to probe Us in vitro (28). However, use of CMCT does not seem feasible for in vivo modification of uridines due to the low membrane permeability of this compound (29). 1-ethyl-3-(3-dimethylaminopropyl) carbodiimide (EDC) has been recently established for uracil probing in different organisms (14,15). However, despite numerous efforts we could not extract RNA from EDC-treated yeast cells because addition of the compound resulted in severe precipitation in the media (data not shown). Therefore, we set out to establish the chemical conditions for use of CMCT for probing uracils in the 5.8S rRNA of *S. cerevisiae*. While EDC effectively modifies several positions on rice 5.8S rRNA (14), none of those sites are reactive with CMCT. However, CMCT demonstrates effective uracil modification in *S. cerevisiae* in vivo at positions U81 and U82 (**Figure 5**). The modification intensity is high at 100 mM CMCT, irrespective of the incubation time. CMCT is too large to effectively penetrate cell wall due to the presence of a quaternary ammonium ion that constitutes a positive charge in CMCT (15), which can explain the weak band intensities due to inefficient entry of CMCT into the cytoplasm (**Figure 5**).

**FIGURE 5.**
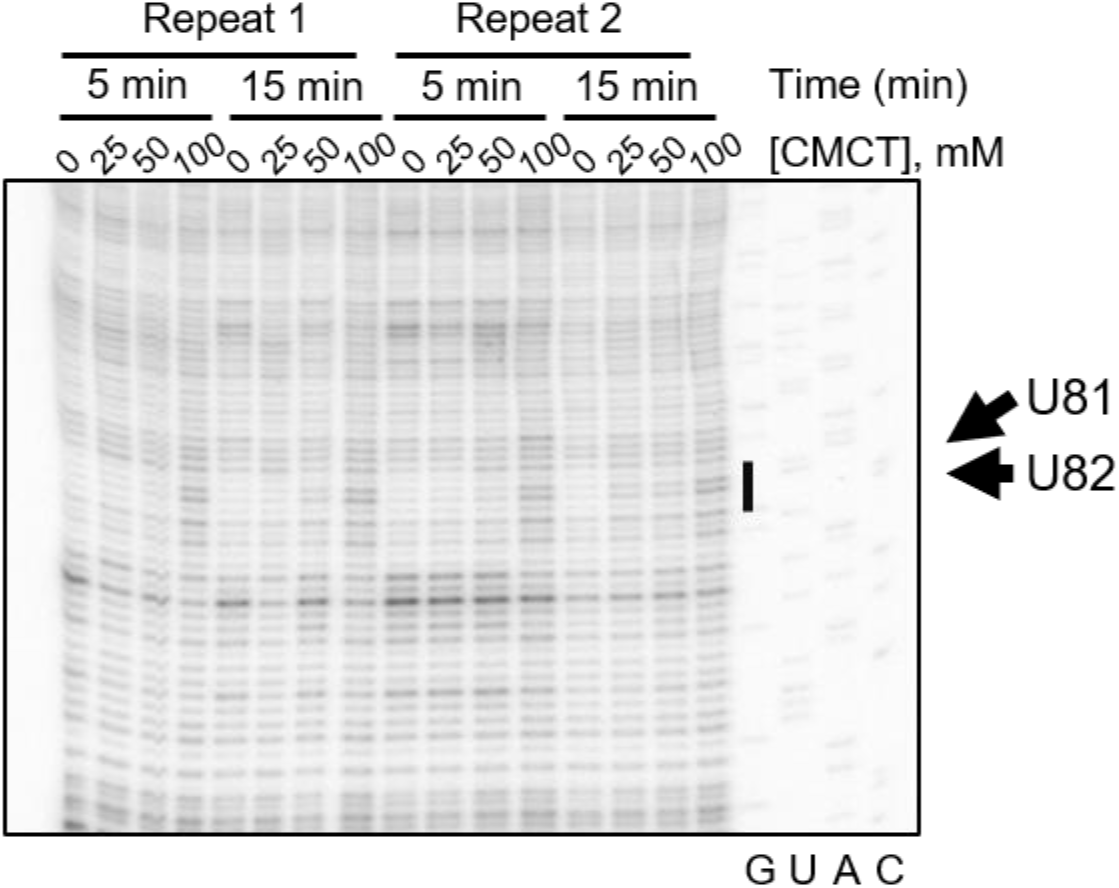
In vivo modification of yeast *S. cerevisiae* 5.8S rRNA by CMCT characterized by denaturing UREA-PAGE of cDNAs after reverse transcription. Reaction conditions at two timepoints and concentrations of glyoxal and its derivatives used are listed. Arrows represent modified RNA nucleotide positions by CMCT.

### Extended CMCT treatment causes RNA processing defects in S. cerevisiae

We next assessed the effect of CMCT treatment on the RNA processing pathways in *S. cerevisiae* using Northern blot (**Figure 6A**). The processing of the 3’ end of U3 snoRNA is not affected in the presence of varying concentrations of CMCT after 5-minute treatment (**Figure 6B**). However, extended treatment with CMCT results in a decrease in the level of U3 precursor (**Figure 6B**). This effect is more pronounced for the pre-tRNA product, the level of which goes down dramatically upon treatment with various concentrations of CMCT after 15 minutes (**Figure 6B**). While a defect in the processing of RNA precursors usually results in the accumulation of the processing intermediates, a decrease in the levels of these intermediates represents either a global destabilization of the RNA species or a defect in the RNA transcription.

**FIGURE 6.**
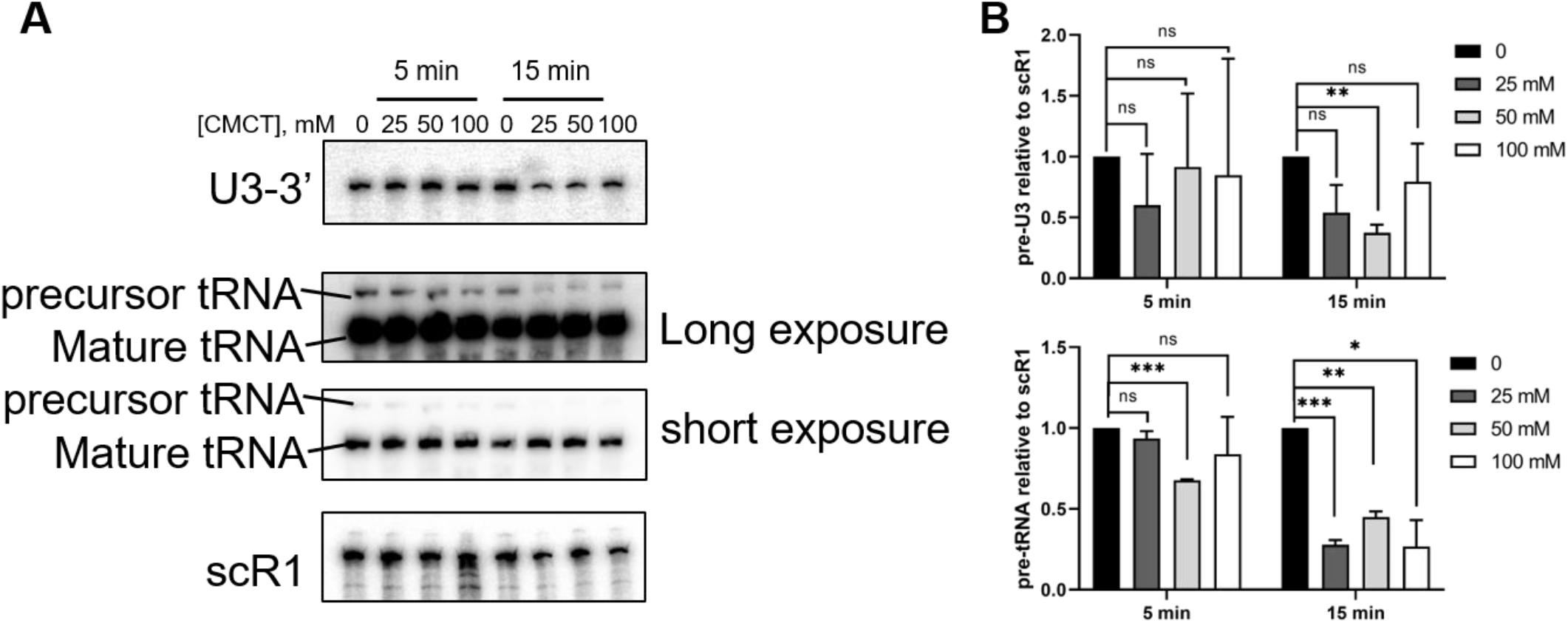
Extensive treatment with CMCT alters the processing of U3 and tRNA^Leu^ in *S. cerevisiae*. **(A)** Northern blot analysis of the U3 and tRNA^Leu^ processing. Reaction conditions at two timepoints are specified at the top of the figure, and concentrations of PGO incubation are listed below reaction time conditions. scR1 RNA is the loading control. **(B)** quantification of the data in panel A. Graphs represent two independent experiments. Significance was determined using a t test; *P ≤ 0.05; **P ≤ 0.01; ***P ≤ 0.001; n.s., nonsignificant.

In summary, this work establishes the optimal chemical conditions in which PGO and CMCT can effectively probe guanine and uracil nucleotides, respectively, in yeast cells. We found that PGO is a potent probe within the glyoxal family derivatives to probe guanine in yeast in vivo. PGO incubation with yeast does not affect its RNA processing pathways, and at the PGO concentrations less than 20 mM and at less than 15-minute incubation period, yeast cells can withstand the toxic effect of PGO. CMCT can be used to probe uracil in yeast in vivo at 100 mM for 5 min, however, proper controls must be done to ensure that it is not interfering with the levels of the RNA of interest.

## Acknowledgement

We thank A.H. Corbett and M. Fasken for helpful discussions. This work was supported by Emory University Research Council (URC) grant to S.K. and NIH grant 1R35GM138123 to H.G.

