## Supplemental Figures for "in vivo RNA structural probing of guanine and uracil nucleotides in yeast"

**Supplementary Figures**

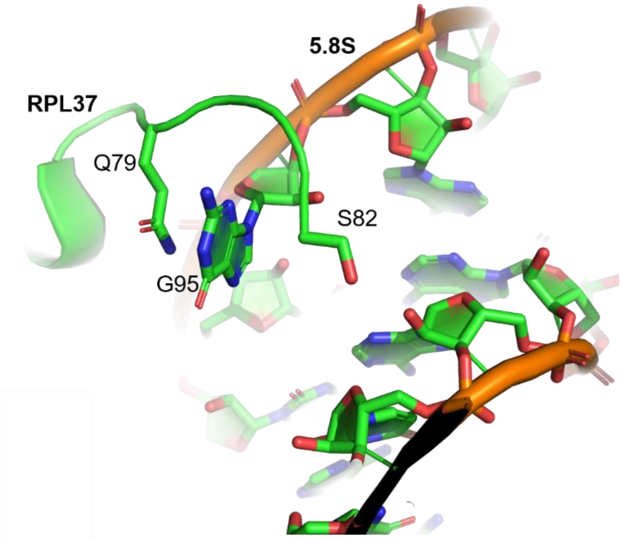


**FIGURE S1. G95 in S. cerevisiae 5.8S is sequestered by the C-terminal tail of RPL37.** Position of 5.8S G95 and RPL37 Gln79 and Ser82 are marked (PDB id: 6TNU). Blue and reg colors represent nitrogen and oxygen, respectively. The phosphate backbone is in orange.

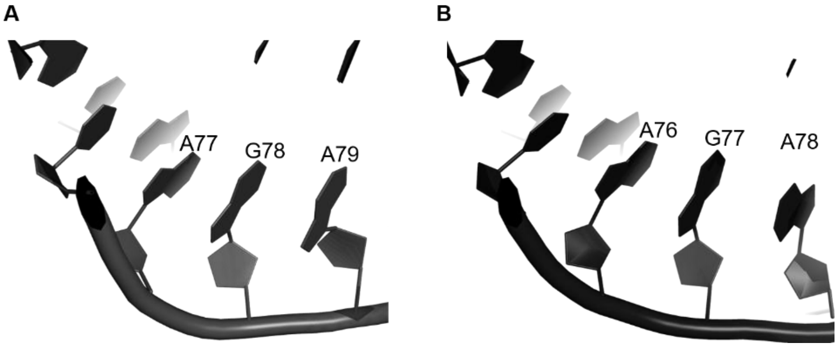


**Figure S2. Structural reason for the lower reactivity of G77 in *C. albicans* 5.8S relative to *S. cerevisiae* G78. (A)** 5.8S from *S. cerevisiae* (PDB id: 6TNU). **(B)** 5.8S from *C. albicans* (PDB id: 6PZY). A79 in *C. albicans* is rotated relative to its counterpart A79 in *S. cerevisiae*.
